# A Spatial Prioritization of Conifer Management to Defend and Grow Sagebrush Cores

**DOI:** 10.1101/2024.02.13.579706

**Authors:** Jason R. Reinhardt, Jeremy D. Maestas, David E. Naugle, Geoffrey Bedrosian, Kevin E. Doherty, Alexander V. Kumar

**Author notes:** **Corresponding author** Reinhardt, J. R., Moscow Forestry Sciences Laboratory, 1221 S. Main St., Moscow, ID 83843, USA.

## Abstract

Sagebrush ecosystems across the western U.S. are in decline due to numerous threats, including expansion of coniferous woodlands and forests. The interagency Sagebrush Conservation Design effort recently quantified sagebrush ecological integrity (SEI) to map remaining core sagebrush areas (relatively intact and functional sagebrush ecosystems) and understand spatial and temporal patterns of change relative to primary threats. This work identified conifer expansion as the second leading cause of decline in sagebrush ecological integrity biome wide. Here, we sought to create a spatial prioritization of conifer management that maximizes return-on-investment to defend and grow core sagebrush areas. Multi-criteria decision analysis (MCDA) was used to incorporate a series of biome-level inputs including SEI, invasive annual grass cover and risk, structural connectivity, and conifer cover and expansion vulnerability into a single prioritization based on collaborative expert input. Our analysis identifies priority areas for conifer management across the sagebrush biome, simulates conifer treatments based on those priorities, and estimates potential changes in SEI as a result of targeted treatment. At a broad scale, we found that the highest priority areas for conifer management were largely located east of the Rocky Mountains. This represents a departure from recent landscape-level trends conifer management efforts in sagebrush systems, which were focused primarily pinyon-juniper expansion in the Great Basin. A majority (52%) of the highest priority areas are managed by the Bureau of Land Management, followed by a large proportion (26%) of priority areas located on privately-owned land – particularly in Wyoming and Montana. Targeting simulated conifer treatments using our prioritization resulted in higher within-core targeting percentages (≥93%) than business-as-usual efforts (23.8%), which would result in a four- to eight-fold reduction in the time to treat priority areas within cores. Finally, we demonstrate that these simulated treatments, targeted with our prioritization, have the capacity to improve SEI in and around treatment areas. This work provides an actionable path to “Defend the Core” as outlined by the Sagebrush Conservation Design effort by helping conservationists more efficiently address conifer expansion in and around core sagebrush areas.

## Introduction

Expansion of coniferous woodlands and forests into adjacent sagebrush (*Artemisia* spp.) rangelands in western North America is a well-documented phenomenon (Romme et al. 2009; Miller et al. 2019; Morford et al. 2024) with a host of undesired consequences for sagebrush-dependent wildlife and ecosystem services (Maestas et al. 2021). In recent decades, the imperiled sage-grouse (*Centrocercus* spp.)—a true sagebrush obligate—has become a primary focus of conifer removal efforts in the region owing to the bird’s sensitivity to tree encroachment (Baruch-Mordo et al. 2013) and favorable response to tree removal (Severson et al. 2017; Olsen et al. 2021; Coates et al., *in Review*). Recent work has demonstrated that the benefits of conifer treatments in sagebrush systems extend to many sagebrush-obligate songbirds as well (Donnelly et al., 2017; Zeller et al., 2021; Tack et al., 2023; Kumar et al., *in Review*). As a result, spatial targeting of conifer tree removal has become a hallmark of sagebrush habitat restoration (Naugle et al., *in Review*), with over half of all treatments clustered within sage-grouse strongholds (Reinhardt et al. 2020). An ever-increasing body of science and geospatial datasets have enabled continual refinement of conservation targeting to better optimize outcomes for sage-grouse (Reinhardt et al. 2017), incorporate multi-species needs (Zeller et al. 2021; Reinhardt et al. 2023; Tack et al. 2023), and improve ecosystem resilience (Chambers et al. 2017). While progress is being made to curtail conifer expansion in some highly targeted landscapes (Naugle et al., *in Review*), range-wide assessments indicate management may just be keeping pace with the rate of expansion (Reinhardt et al. 2020) and conifer encroachment remains the second leading driver of sagebrush ecosystem degradation (Mozelewski et al., *in Review*) indicating room for further improvement strategic conservation.

The Sagebrush Conservation Design (SCD; Doherty et al. 2022) represents the latest evolution of strategic conservation planning in the sagebrush biome, providing a comprehensive and spatially explicit view of sagebrush ecosystem threats and change through time. The SCD seeks to identify intact and functioning sagebrush ecosystems relative to primary threats to enable a more proactive, ecosystem-based conservation strategy. Using remotely-sensed data, the SCD models Sagebrush Ecological Integrity (SEI) across sagebrush rangelands as a function of sagebrush cover, perennial grass cover, annual grass cover, human modification, and tree cover (Doherty et al., 2022) in an effort to identify Core Sagebrush Areas (CSAs; most intact), Growth Opportunity Areas (GOAs; moderately intact), and Other Rangeland Areas (ORAs; least intact). The SCD encourages managers to implement a new conservation philosophy: “Defend the Core, Grow the Core, Mitigate Impacts” (NRCS, 2021), which emphasizes prioritizing the protection and management of intact “core” sagebrush ecosystems over restoration of more highly degraded areas. This approach – referred to here as Defend the Core – is designed to direct resources and management efforts first into priority landscapes where early intervention to address threats is more likely to be effective and sustainable long term.

Here, we leverage the SCD to improve spatial prioritization of conifer management across the sagebrush biome to help land managers better maintain and improve ecological integrity of sagebrush ecosystems in the face of rapid change. Using a collaboratively-informed modeling approach, we developed a biome-wide prioritization for conifer management that incorporates SEI, conifer cover and expansion risk, invasive annual grass risk, and ecosystem-level connectivity. Our goals were to: 1) create a biome-wide prioritization layer that can be used to enhance outcomes of future conservation investments, 2) demonstrate the utility of expert-informed modeling to address conifer expansion, and 3) evaluate how our prioritization can improve landscape-scale ecological integrity.

## Methods

### Study area

This study considers the full extent (99.1 million hectares) of the sagebrush biome in the United States (Jeffries and Finn, 2019). Non-rangeland areas were masked out from the analyses (Maestas et al., 2020), as were alpine tundra (USGS, 2016). Land cover in the study area primarily consists of sagebrush ecosystems, numerous types of pinyon-juniper woodlands, and agricultural land (Miller et al., 2008; Homer et al., 2015). The climate across the sagebrush biome tends to be cold and semiarid with hot and dry summers but cold and wet winters (Kottek et al., 2006). There is, however, significant variation across the wide latitudinal gradient; the climate in the Southern Great Basin is particularly different, with warmer summers and less overall precipitation (Kottek et al., 2006; Daly et al., 2008).

### Data

Estimated conifer cover was obtained by taking the tree cover dataset from the Rangeland Analysis Platform (v3, Allred et al., 2021) and masking it to the sagebrush biome as described above. This dataset provides coverage over our entire study area and has a mean absolute error of 2.6% (Allred et al., 2021). A conifer expansion (i.e., dispersal and establishment) risk layer was produced using a 200-m dispersal kernel centered on pixels with conifer cover >2% (to account for error and the possibility of misidentifying trees where there are none). A distance of 200-m was used based on a rounded-up estimate of the fastest rate of pinyon pine expansion computed by Vander Wall (2023). Areas further from existing conifer along this gradient were considered to be at lower risk for conifer expansion.

To support proactive management, we wanted our prioritization to preferentially target the leading edge of conifer expansion with little or no conifer cover but with the risk of conifer expansion, followed by areas of low conifer cover, with priority gradually decreasing as cover increases. To accomplish this, we created a single conifer component by combining the conifer cover dataset with the conifer expansion risk layer. This was done by truncating the conifer cover layer at 2% (matching our dispersal kernel threshold, above) and inserting a rescaled conifer expansion risk in the 0-2 values of the cover dataset. This put the highest emphasis for our prioritization analyses on areas of expansion risk and low cover (Phase I *sensu* Miller et al., 2005) woodlands and forests. At the ground level, this means that the priority would be tilted toward treating Phase I (Miller et al., 2005) pinyon-juniper woodlands and preventative or early intervention treatments in shrub systems on the leading edge of conifer expansion.

A quantification of SEI (Doherty et al., 2022) computed without the conifer threat component was used in these analyses to prioritize areas which demonstrate high ecological integrity in the absence of conifer cover. We incorporated the relative risk of 5 invasive annual grasses (cheatgrass (*Bromus tectorum*), red brome (*Bromus rubens*), Japanese brome (*Bromus japonicus*), ventenata (*Ventenata dubia*), and medusahead (*Taeniatherum caput-medusae*)) based on data from Boyd et al. (*in Review*), which represents the product of current invasive annual grass cover (average from 2019-2022) derived from the Rangeland Analysis Platform (Allred et al., 2021) as well as modelled suitability for invasive annual grasses (Jarnevich et al. 2023a, Jarnevich et al. 2023b, Williams et al. 2023). Landscape-level structural connectivity derived from a multi-scale model using a resistant Gaussian kernel approach (produced by Theobald et al., *in Review*) was incorporated to represent broad ecosystem-level connectivity across the sagebrush biome.

### Analyses

We used multi-criteria decision analysis (MCDA; Belton and Stewart, 2002; Marttunen et al., 2017) via the analytical hierarchy process (Saaty, 1987) to develop a spatial prioritization of conifer management across the sagebrush biome. A collection of collaborative subject matter experts (*n* = 19) in fields including rangeland ecology, landscape ecology, invasive plant management, wildlife ecology, conservation biology, and forestry were asked to independently rank the pairwise importance (*sensu* Saaty, 1987) of each of the 4 criteria included in this analysis: conifer, SEI, invasive annual grass risk, and structural connectivity. These experts were drawn from the broader Sagebrush Conservation Design group. Responses were transformed into a summary matrix and the right eigenvector was computed using the pairwiseComparisonMatrix() and calculateWeights() functions in the *FuzzyAHP* R package (Caha and Drážná, 2019; R Core Team, 2023). These computed weights were used to combine the standardized criteria datasets into a single conifer prioritization map representing priority on a 0-100 scale. In computing the MCDA, the directionality of each input dataset was configured such that the highest values would represent areas of relatively low conifer threat, low invasive annual grass presence and risk, high SEI, and high structural connectivity. To evaluate the distribution of priority areas across land tenure, we split the prioritization dataset into deciles and summarized across ownerships. We evaluated the distribution of the top 3 priority deciles (8^th^ – 10^th^) across ownerships as described in the BLM Surface Management Agencies database (BLM, 2023).

To illustrate the utility of this prioritization, we simulated conifer treatments in several different scenarios. We simulated targeted conifer treatments by using an area estimate of business-as-usual (BAU) annual conifer treatment (64,319 ha) computed by Mozelewski et al. (*in Review*). We selected an area equal to this from the highest priority locations as estimated by the MCDA prioritization. To evaluate the potential added gain in ecological integrity by increasing treatment area, we doubled the BAU area in a second set of simulated conifer treatments (MCDA x2). We also evaluated the potential distribution of 10 years of conifer treatments for both the BAU (10 y MCDA) and double-area (10 y MCDA x2) scenarios, resulting in a total of 4 conifer treatment scenarios.

Mozelewski et al. (*in Review*) quantified the distribution of contemporary conifer treatments occurring within each ecological integrity category (ORAs, GOAs, and CSAs) and estimated that 23.8% of contemporary conifer treatments currently occur within CSAs. To compare potential differences in targeting between MCDA scenarios and BAU, we evaluated the proportion of our simulated conifer treatments that occurred within CSAs for each MCDA. We then compared the relative amount of within-core treatment area required to completely treat priority areas within cores. The ‘priority areas’ used for this analysis consisted of the top 50% of the MCDA-derived priority scale, and the ‘highest priority’ areas consisted of the top 25%. Within-core acreage of these priority categories was computed, and then compared with the within-core targeted treatments for the BAU and doubled area treatment scenarios along with numbers from the current and most optimistic conservation scenarios outlined by Mozelewski et al (*in Review*).

The potential impact of simulated conifer treatments on SEI was evaluated by setting target areas to zero conifer cover for each scenario and recomputing SEI as per Doherty et al. (2022). Treatments could potentially have impacts on ecological integrity beyond their immediate area of simulated implementation, as the ecological integrity model is computed at a broader spatial scale (560m vs 30m) using a Gaussian kernel (Doherty et al., 2022); this also means, however, that treatments have the potential to be “washed out” in the moving-window smoothing process if they are not large enough. Changes in ecological integrity due to simulated treatments were summarized by category: ORAs, GOAs, and CSAs. The area of change was summarized in terms of whether a particular area remained stable within a category, demonstrated an increase of ecological integrity within a category, or improved to the next category up (e.g., GOA to CSA). Finally, we evaluated both the distribution and the potential impact of simulated conifer treatments across land tenure categories as found within the BLM Surface Management Agencies database (BLM, 2023).

## Results

The MCDA weights produced via AHP put SEI first (0.525), followed by both conifer and invasive annual grass (0.170) and then connectivity (0.135) (Table 1). The prioritization based on collaborative input therefore puts the most weight on SEI (without the conifer threat component), equal weight on conifer and invasive annual grasses, followed closely by connectivity. High priority areas are thus reflective of locations with relatively high ecological integrity, low invasive annual grass presence and risk, low conifer threat, and higher landscape-level connectivity.

**Table 1.** Criteria weighting as derived from the analytical hierarchy process (Saaty, 1987). Collaborative responses represent input from a range of natural resource experts (*n* = 19).

| Criterion | Weight |
| --- | --- |
| Conifer | 0.170 |
| Ecological Integrity | 0.525 |
| Invasive Annual Grass | 0.170 |
| Connectivity | 0.135 |

The largest portion of the highest priority deciles (8-10) in our prioritization fall within land managed by the Bureau of Land Management (BLM; 52%), followed by private ownership (26%) (Tables 2-4). Approximately 80% of the high priority areas occurring on land managed by the BLM occurs within three states: Idaho, Montana, and Wyoming (Table 3). Similarly, the distribution of privately-owned high priority areas varies greatly by state (Table 4), with private landowners in Wyoming (48%) and Montana (37%) making up the bulk of this category. While State agencies only manage 11% of the total high priority areas, the amount of this area in the 9^th^ and 10^th^ deciles is relatively high (∼19% and ∼40% of the area in each decile, respectively), suggesting that state agencies manage a relatively high proportion of areas with the highest MCDA prioritization ratings (Table 2).

**Table 2.** Area distribution of the highest priority areas for conifer management as identified by MCDA across major ownership groups. The top 3 deciles of priority (8-10) are shown.

| Area (ha) by Priority Decile |  |  |  |  |  |
| --- | --- | --- | --- | --- | --- |
| Ownership | 8 | 9 | 10 | Top 3 Deciles | Percentage |
| BLM | 764063 | 183761 | 9021 | 956845 | 52% |
| Private | 420373 | 61751 | 2991 | 485115 | 26% |
| State Agencies | 128533 | 62856 | 8239 | 199629 | 11% |
| USFS | 119803 | 17936 | 109 | 137847 | 7% |
| Tribal/BIA | 24052 | 3648 | 0 | 27700 | 2% |
| Other Public | 30421 | 3729 | 30 | 34180 | 2% |
| **Total** | **1487245** | **333682** | **20390** | **1841317** |  |

**Table 3.** Area distribution across states of the highest priority areas for conifer management as identified by MCDA within land managed by the BLM. The top 3 deciles of priority (8-10) are shown.

| Area (ha) by Priority Decile |  |  |  |  |  |
| --- | --- | --- | --- | --- | --- |
| State | 8 | 9 | 10 | Top 3 Deciles | Percentage |
| AZ | 952 | 0 | 0 | 952 | 0% |
| CA | 11106 | 657 | 0 | 11763 | 1% |
| CO | 19997 | 424 | 0 | 20421 | 2% |
| ID | 109830 | 47677 | 6683 | 164190 | 17% |
| MT | 128380 | 34790 | 20 | 163191 | 17% |
| NM | 213 | 0 | 0 | 213 | 0% |
| NV | 53620 | 2507 | 0 | 56127 | 6% |
| OR | 61002 | 1906 | 0 | 62908 | 7% |
| UT | 27981 | 11033 | 1605 | 40619 | 4% |
| WY | 350981 | 84766 | 713 | 436461 | 46% |
| **Total** | **764063** | **183761** | **9021** | **956845** |  |

**Table 4.** Area distribution across states of the highest priority areas for conifer management as identified by MCDA within the Private ownership group. The top 3 deciles of priority (8-10) are shown.

| State | Area (ha) by Priority Decile |  |  | Top 3 Deciles | Percentage |
| --- | --- | --- | --- | --- | --- |
|  | 8 | 9 | 10 |  |  |
| AZ | 2 | 0 | 0 | 2 | 0% |
| CA | 177 | 0 | 0 | 177 | 0% |
| CO | 2311 | 19 | 0 | 2330 | 0% |
| ID | 21135 | 7168 | 2618 | 30922 | 6% |
| MT | 156718 | 23078 | 303 | 180098 | 37% |
| NM | 632 | 0 | 0 | 632 | 0% |
| NV | 7254 | 580 | 0 | 7834 | 2% |
| OR | 3532 | 30 | 0 | 3562 | 1% |
| UT | 24178 | 3535 | 65 | 27779 | 6% |
| WY | 204434 | 27340 | 5 | 231779 | 48% |
| **Total** | **420373** | **61751** | **2991** | **485115** |  |

Simulated treatment area was a function of the MCDA scenario used (Table 5), with one year of MCDA-targeted treatments encompassing ∼57,500 ha, up to 10 years of treatments using the MCDA x2 scenario covering ∼1,332,000 ha. The BLM held a plurality of simulated treatment areas; 50-55% of treated areas were found on BLM-managed land across scenarios (Table 5). State agencies demonstrated relatively high proportions (∼37% and ∼29%) of treatment in the single-year treatment scenarios relative to the 10 year scenarios (∼16% and ∼12%) (Table 5). Private landowners were allotted more area in the 10 year scenarios (∼21% and ∼24%) than the single-year scenarios (∼11% and ∼12%) (Table 5). These contrasts reflect differences in available high priority areas for treatment targeting across ownership (Table 2).

**Table 5.** Area distribution of simulated treatment area across ownerships, by treatment scenario.

| Ownership | Treatment Area (ha) by Scenario |  |  |  |
| --- | --- | --- | --- | --- |
|  | MCDA | MCDA x2 | MCDA 10 yr | MCDA x2 10 yr |
| BLM | 28502 | 71123 | 348314 | 704037 |
| Private | 6523 | 15816 | 137823 | 331028 |
| State | 21344 | 37549 | 99664 | 158214 |
| USFS | 780 | 3536 | 40441 | 97269 |
| BIA/Tribal | 0 | 123 | 7637 | 18524 |
| Other Public | 405 | 1028 | 8897 | 22998 |
| **Total** | **57554** | **129174** | **642777** | **1332070** |

Treatment scenarios resulted in within-core targeting ranging from 93.01% (10 y BAU x2) to 96.59% (BAU) (Table 6). Within-core targeting percentage decreased as the amount of simulated treatment area increased across scenarios (i.e., from BAU to 10 y BAU x2); the single-year treatment scenarios had slightly higher within-core percentages (96.49% and 96.41%) than the 10 year scenarios (94.84% and 93.01%), as a result of more GOA area being selected as more total area was available for treatment.

**Table 6.**
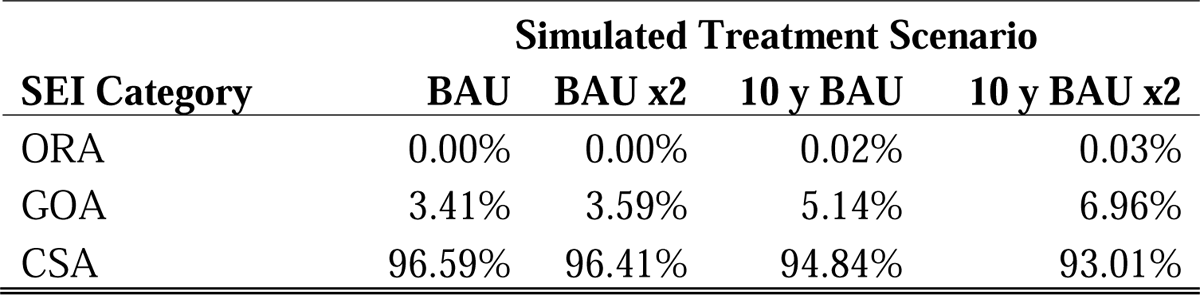
Distribution of simulated treatment area across SEI categories, by treatment scenario.

| SEI Category | Simulated Treatment Scenario |  |  |  |
| --- | --- | --- | --- | --- |
|  | BAU | BAU x2 | 10 y BAU | 10 y BAU x2 |
| ORA | 0.00% | 0.00% | 0.02% | 0.03% |
| GOA | 3.41% | 3.59% | 5.14% | 6.96% |
| CSA | 96.59% | 96.41% | 94.84% | 93.01% |

The implications of this relatively high rate of within-core targeting was explored by comparing targeting percentages with the total area of ‘priority’ and ‘high priority’ landscapes within CSAs (top 50% of priority and top 25% of priority, respectively). We estimate that at BAU treatment acreages, a prioritization-informed targeting strategy would take approximately 19 years to treat the highest priority within-core areas, and 143 years to treat all priority areas (Table 7). These numbers are nearly halved when doubling the BAU acreage, and this particular scenario (MCDA x2) is relatively close to the most optimistic conservation scenario outlined by Mozelewski et al. (*in Review*) (Table 7). Both prioritization-informed scenarios have a demonstrably shorter time-to-treat than the ‘Current’ scenario, which is based on a proportion of 23.8% of treatment areas within CSAs (Mozelewski et al., *in Review*).

**Table 7.** Estimated years to treat all available area within each category, by scenario. “Current” is based on the current rate of treatment and distribution within cores (23.8%) as per Mozelewski et al. (*in Review*); “MCDA” is based on the same acreage but using the prioritization-informed targeting of 96.59% of treatment areas within cores; “MCDA (double)” represents the 2x acreage scenario with a prioritization-informed targeting of 96.41% in cores; and “Strategic All-In” represents the most optimistic conservation scenario outlined by Mozelewski et al. (*in Review*), with 80% of treatments within cores.

| Scenario | Years to Treat |  |
| --- | --- | --- |
|  | Core: Highest | Core: All Priority |
| Current | 78.19 | 579.12 |
| MCDA | 19.30 | 142.94 |
| MCDA (double) | 9.67 | 71.60 |
| Strategic All-In | 10.87 | 80.51 |

Simulated treatments resulted in an increase in SEI (Table 8, Figure 3). The amount of SEI gain was roughly correspondent to the area treated, with the single year treatment scenarios demonstrating lower gain than the 10-year scenarios, and the BAU scenarios lower than the doubled acreage scenarios (Table 8). The relative distribution of SEI gain across SEI categories as outlined in Table 8 remained relatively stable as the amount of treatment area increased (BAU to 10 y BAU x2).

**Table 8.**
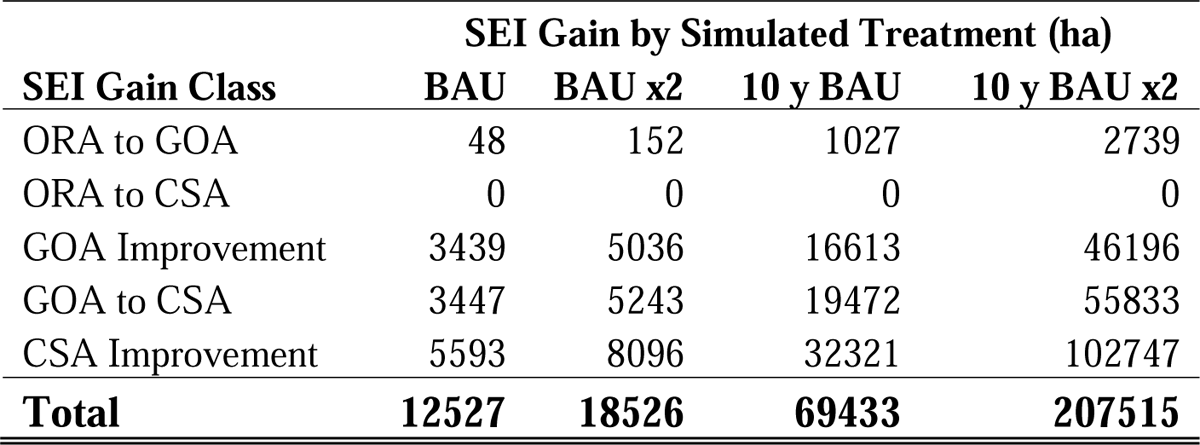
Summary of the study-wide area of gain in SEI as a result of simulated conifer treatments, by scenario.

## Discussion

We produced a collaboratively informed prioritization for conifer management to defend and grow CSAs across the sagebrush biome (Fig. 1). Given the vast extent of the biome (99.1 M ha), large coverage of even the CSAs (13.5 M ha; Doherty et al., 2022), rate of conifer expansion (Filippelli et al., 2020), and current pace of conifer management (Mozelewski et al., *in Review*), better spatial targeting of limited resources is clearly needed to address second leading cause of degradation in sagebrush ecological integrity. Our work shows defending and growing CSAs is possible with shifts in biome-wide prioritization of conifer management. This may require forging new partnerships across private and public lands in regions where “conservation readiness” (Wollstein et al., *in Review*) for conifer management at landscape scales may not exist and needs to be built.

**Figure 1.**
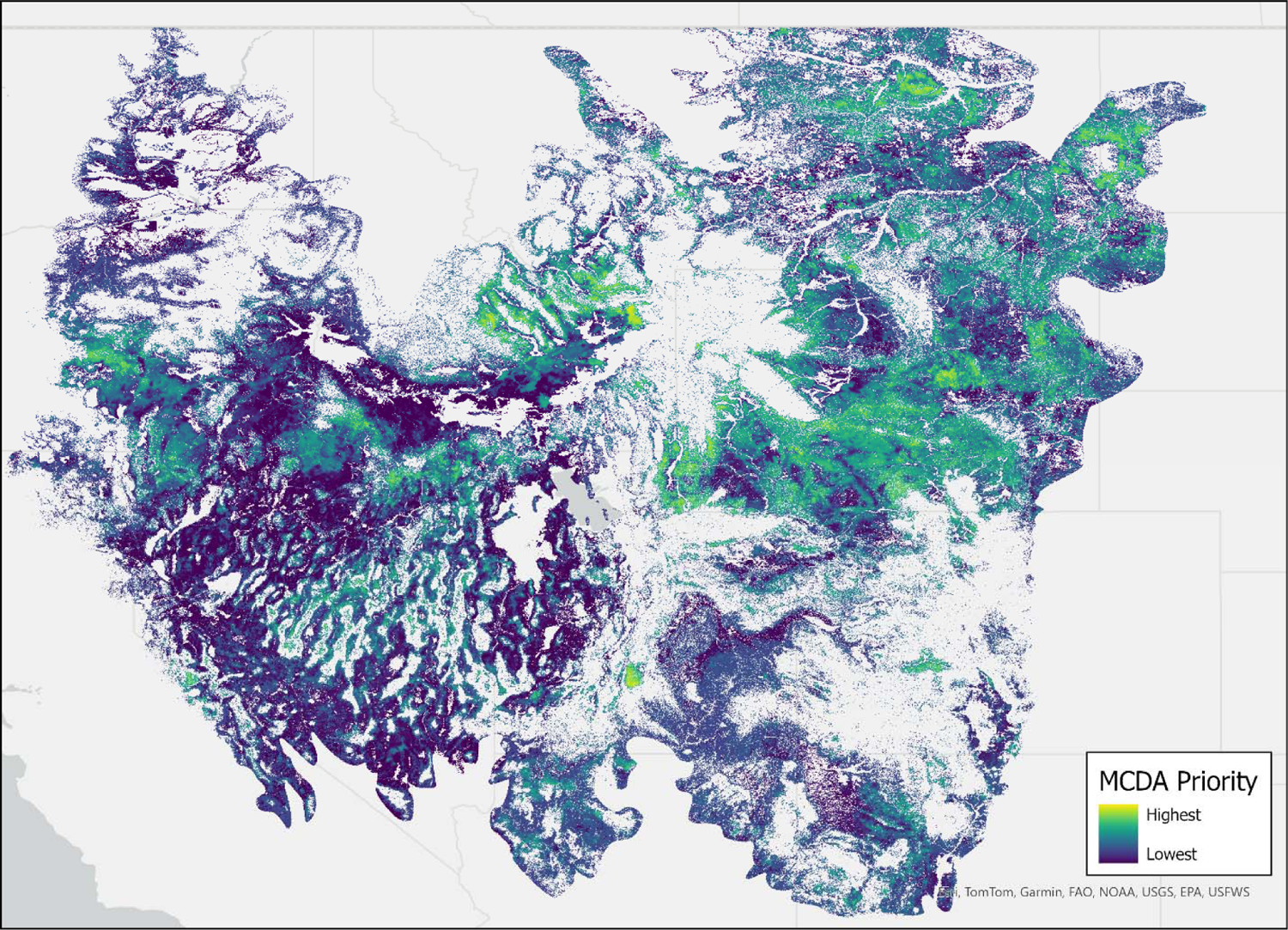
Conifer management priority as computed by MCDA (30-m resolution). Scale is unitless and represents relative priority. Yellow to green represent priority areas, with darker colors representing areas of decreasing priority. Non-rangeland and alpine areas are masked (USGS, 2016; Maestas et al., 2020)

Among the most notable patterns emerging from our prioritization is the location of the highest priority areas for conifer management (Fig. 2, Tables 2-4). Many of the top priority areas identified are located east of the Rocky Mountains, contrasting with the broad-scale efforts of sagebrush conservation for much of the 2010s which emphasized priority landscapes primarily within the Great Basin (Naugle et al., *in Review*). This outcome stems from the model inputs and weights prioritizing opportunities to address early conifer expansion in areas of high ecological integrity, with higher connectivity, and least impacted by invasive annual grasses (Table 1). The large-scale spatial shift suggests that targeted conifer management in the eastern portion of the biome may provide more “low-hanging fruit” opportunities today to help achieve SCD goals of defending and growing CSAs. However, our work also indicates continued opportunities for improved targeting in the Great Basin, the need for maintenance of previous treatments, and the importance of coupling conifer management with invasive grass control (Naugle et al., *in Review*; Smith et al., *in Review*).

**Figure 2.**
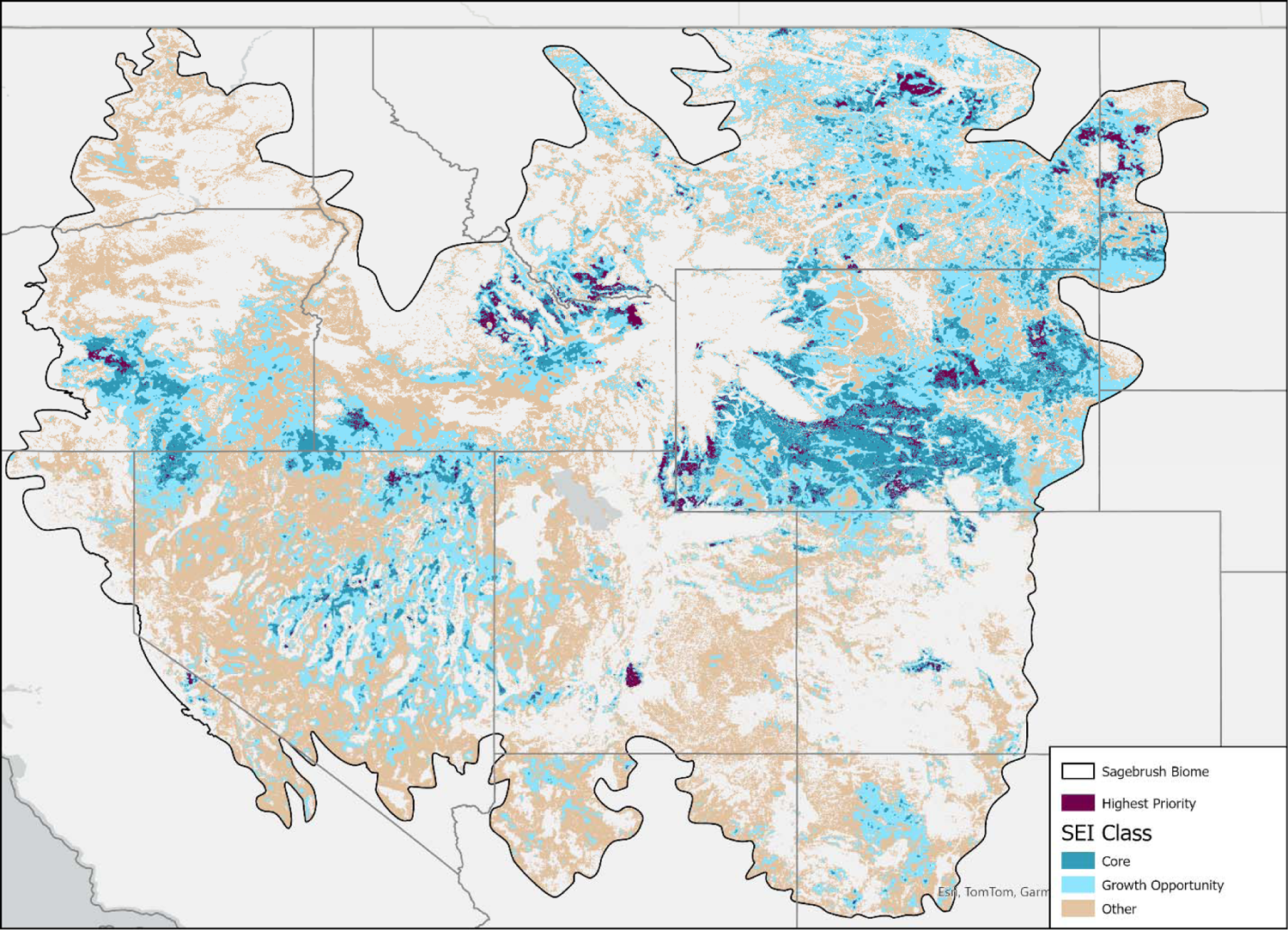
Highest priority (top 3 deciles) conifer management areas (purple), as estimated by the MCDA prioritization. SEI classes shown in dark blue (CSA), light blue (GOA), and tan (ORA).

**Figure 3.**
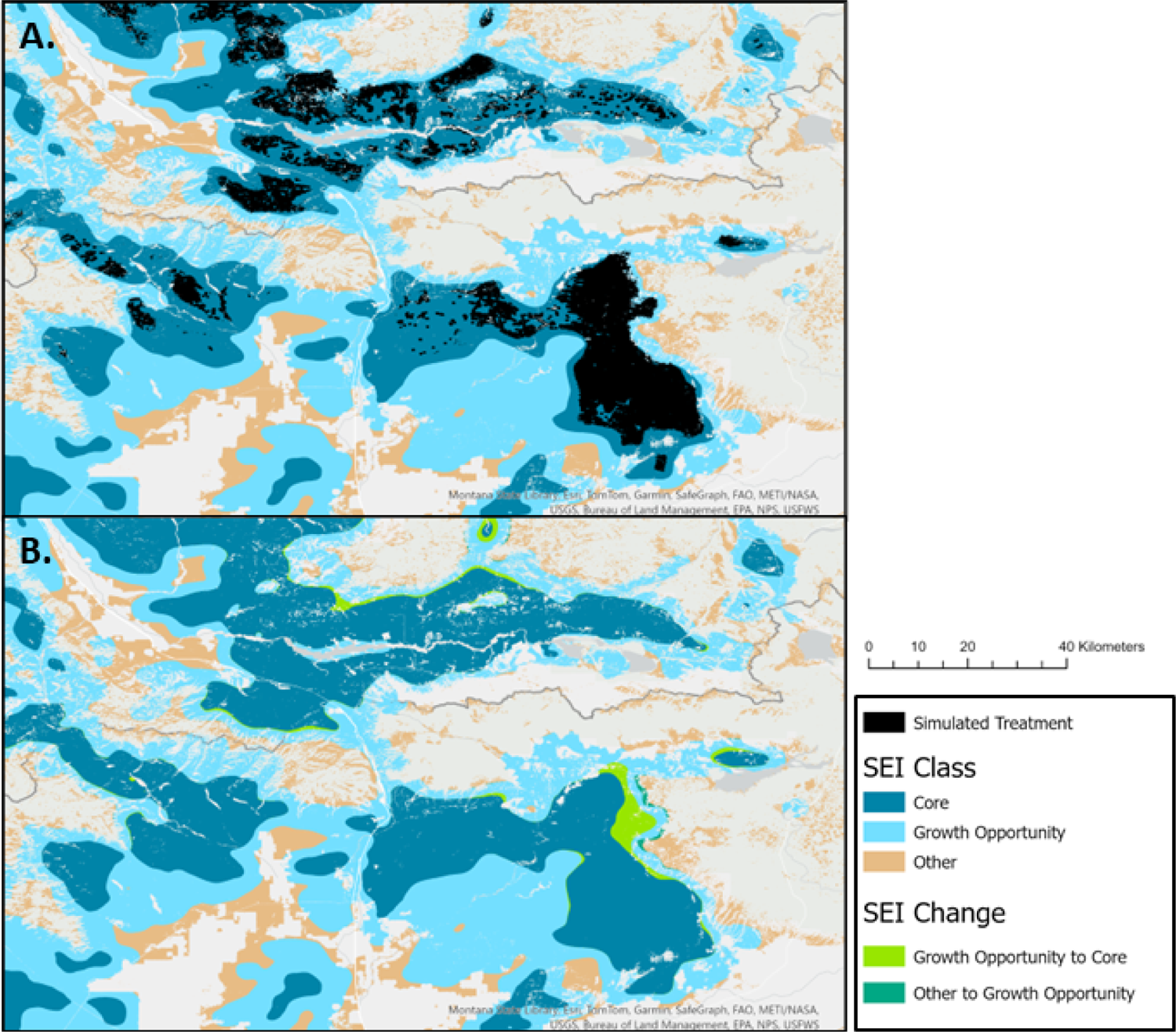
SEI classes and locations of simulated conifer treatments for 10 years of BAU acreage (A.) and SEI improvements as a result of simulated treatments (B.). The focal area is eastern Idaho and southwestern Montana, an area of especially high concentration of high priority areas for conifer management.

BLM and private lands accounted for 78% of the highest priority areas identified for conifer management (Table 2). The distribution of these high priority areas across public and private lands reemphasizes the need to work across ownerships to meet conservation goals (Wollstein et al., *in Review*). BLM priority areas were primarily located in Wyoming, Idaho, and Montana (Table 3), whereas private land opportunities were heavily concentrated in Montana and Wyoming (Table 4). Previous work has demonstrated the urgency of the issue in Montana where rangelands are being converted into woodlands and forests at a rate exceeding 58,000 ha/yr (Morford et al., 2024). Local examples of cross-boundary collaboration to address this threat are already emerging in communities like southwestern Montana, providing a model for others in these states as they seek to scale up management (see: https://www.partnersinthesage.com/sw-montana-geography-of-sagebrush). The Natural Resources Conservation Service’s Working Lands for Wildlife partnership is also supporting conifer removal east of the Rocky Mountains through technology transfer, training, and program funding (Naugle et al., *in Review*).

Lands managed by the BLM possessed the plurality of simulated treatment acreage across scenarios (Table 5), though there were differences in the relative proportions of treatment areas when moving up from single-year to 10 year scenarios; in the former, state agencies made up a relatively large portion of available treatment area, while in the latter private ownership and the USFS demonstrated a higher proportional area. This reflects the differences in abundance of the highest priority areas available for treatment; state agencies manage a relatively large portion of the highest priority areas on the landscape (Table 2), which were preferentially targeted first in the single-year treatment scenarios.

Mozelewski et al. (*in Review*) estimate approximately 64,319 hectares of conifer management occurring per year in support of sagebrush ecosystem conservation, with an estimated 23.8% of those actions occurring within CSAs. Our simulated conifer treatments, targeted using our prioritization, demonstrated a much higher rate of within-core ‘management’, ranging from 93.01% to 96.59% depending on area targeted (Table 6). Given that ecological integrity was a highly weighted criterion in our analysis (Table 1), this was both expected and a desirable outcome. Indeed, Naugle et al. (*in Review*) found a 20% increase in effectiveness when treatments were saturated in a “Defend the Core” strategy, highlighting the importance of deliberately targeting treatments within CSAs. With this in mind, we explored the potential treatment timeline impacts of our prioritizations. Relative to the BAU targeting regime of 23.8%, we found that the prioritization-targeted treatments using both BAU and doubled treatment areas led to a four- to eight-fold drop in the estimated years to treat currently existing priority habitat within cores (78.19 years to 19.30 (BAU) and 9.67 (BAU x2) years; Table 7), the latter even surpassing the most optimistic conservation scenario (10.87 years) outlined by Mozelewski et al. (*in Review*).

While this prioritization targeting is demonstrably effective at increasing within-core treatment area, the overall within-core area is quite large (13.5 M ha) and the highest priority areas are only a subset of this. Managing the full available area within cores, whether for conifer removal, prevention, or monitoring would take longer, and this is evident in the temporal gulf between treating only the highest priority areas (19.3 years) or all priority areas (142.94 years) – the difference is nearly an order of magnitude (Table 7). The highest priority areas characterize areas that not only fall within CSAs but are desirable for treatment based on the other criteria in the analysis: they demonstrate lower conifer cover, less risk from invasive annuals both prior to and post-treatment, and represent high landscape-level connectivity. In the absence of abundant resources to pursue sagebrush conservation efforts, these are factors that should be considered when planning conifer treatments (Roundy et al., 2014; Bybee et al., 2016; Fick et al., 2022; Boyd et al., *in Review*; Naugle et al., *in Review*; Theobald et al., *in Review*).

The prioritization-targeted treatments simulated here resulted in improvements in SEI (Table 8; Figure 3), and the relative amount of improvement was broadly a function of area treated. Given the highly targeted nature of the simulated treatments, it follows that the majority of improvement in SEI was within CSAs and GOAs, with little improvement in ORAs. This intentional targeting supports the Defend the Core strategy and has the potential to realize benefits from saturating treatments within priority areas (Naugle et al., *in Review*; Figure 3). The overall area of SEI improvement was less than the total area treated. There are two major reasons for this: first, the calculation of SEI is performed using a gaussian kernel at a larger spatial grain than either the prioritization or simulated treatment datasets (560m vs. 30m), so simulated treatment areas must be large enough to be adequately represented in the moving window. Second, SEI is computed based on several factors other than conifer cover, including sagebrush cover, perennial herbaceous cover, invasive annual grasses, and human modification (Doherty et al., 2022), so it is possible that even a reduction in conifer threat would not be enough to meaningfully increase SEI for a given area.

At the biome level, conifer expansion remains an ongoing threat to sagebrush ecosystems (Maestas et al., 2021). It is important, however, to note that there is a layer of complexity with respect to conifer expansion into rangeland systems. Recent work suggests that there may be impacts of drought and climate change on some pinyon pine and juniper species in some areas (Snyder et al., 2019; Shriver et al., 2022; Noel et al., 2023), but contemporary remote sensing data clearly illustrate conifer expansion in all but the southernmost ecoregions of woodland distribution (Filippelli et al., 2020; Allred et al., 2021; Kleinhesselink et al. 2023; Morford et al. 2024). This apparent contradiction may be a result of complex site and stand dynamics interacting with recent climatic changes at multiple scales (e.g., Flake and Weisberg, 2019; Schultz et al., 2022; Schultz et al., 2023). Future prioritization efforts, especially those which seek to project the changing spatial distribution of priority conifer management areas over time, should carefully consider these complex interactions – such efforts may be best served by creating separate prioritization models for different ecoregions.

Growing conservation concern around pinyon jay (*Gymnorhinus cyanocephalus*) is another emerging challenge when considering how to prioritize conifer management efforts in support of sagebrush conservation (Boone et al., 2018; Zeller et al., 2021; Reinhardt et al., 2023). Our prioritization did not explicitly include pinyon jay as a factor, but the areas we identify as high priority (Figure 2) have little overlap with pinyon jay strongholds modeled using Breeding Bird Survey data (Tack et al., 2023). Local and regional context are essential when considering potential nontarget effects of conifer management to support sagebrush ecosystem management. At smaller spatial extents where management decisions are geographically confined, such as counties, BLM field districts, and other federal properties (Doherty et al., *in Review*), decisionmakers – particularly in central Nevada or Utah where priority areas for conifer management may intersect with pinyon jay strongholds (Tack et al., 2023) — managers may wish to consider a holistic conifer management approach that includes restoration objectives tailored specifically to sagebrush rangelands and pinyon-juniper woodlands or pinyon jay habitats (Maestas et al. 2021).

## Implications

Our work builds upon an evolving history of strategic conifer management across the sagebrush biome over the last two decades (Reinhardt et al., 2023) by stepping up the focus from individual species to the ecosystem itself (Doherty et al., 2022). The prioritization was created with collaborative input from experts across natural resource fields and represents a framework for targeting limited resources to address conifer expansion. The highest priority areas identified here directly support the Defend the Core strategy, as illustrated by the placement and impact of simulated conifer treatments targeted using the prioritization. Deliberately targeted treatments in areas with low conifer cover and low risk of annual grass invasion represent the highest value for conservation resources directed at conifer management, and this tactic is further strengthened by saturating treatments within high priority landscapes (Naugle et al., *in Review*) and across ownership boundaries (Smith et al., *in Review*).

Our prioritization of conifer treatments in the sagebrush biome provides a tool for resource managers to strategically target treatment areas that are more likely to be effective at improving landscape-scale rangeland conditions, as measured here using SEI. The scale and impact of past conservation efforts has been insufficient to halt the spread of conifer encroachment and associated loss of sagebrush rangelands (Mozelewski et al., *in Review*) in all but a few landscapes (Naugle et al., *in Review*). Strategic prioritization of conservation efforts will be required to retain the highest quality sagebrush cores, even under scenarios accounting for substantial increases in conservation volume. By leveraging a collaboratively informed model that includes multiple conservation targets, our work represents an opportunity to improve how the conservation community delivers treatments from biome-wide to regional scales.

Recent sagebrush restoration actions targeting conifers have largely focused efforts on the Great Basin (Reinhardt et al., 2020; Mozelewski et al., *in Review*; Naugle et al., *in Review*), but our prioritization identified the largest area of highest priorities in the eastern and northern portions of the sagebrush biome (Fig. 2). This suggests that the “low hanging fruit” opportunities today are located in areas characterized by high SEI and early-stage conifer encroachment, as opposed to working in landscapes impacted by multiple threats where treatment costs are higher and success is limited. Our prioritization can be used alongside multiple complementary targeting tools (Bedrosian et al., *in Review*; Boyd et al., *in Review*; Cross et al., *in Review*; Holdridge et al., *in Review*, Theobald et al., *in Review*) to support an ecosystem-based approach to defending and growing intact sagebrush rangelands.

## Acknowledgements

We thank the Sagebrush Conservation Design collaborative group for their expert input used in our analysis, and for providing context and inputs for our analysis; special thanks to Dave Theobald and Tina Mozelewski for the latter. This research was supported by the USDA Forest Service, Rocky Mountain Research Station. The findings and conclusions presented in this paper are those of the authors and do not necessarily reflect the views of the USDA Forest Service, US Fish and Wildlife Service, or USDA Natural Resources Conservation Service.

## Declaration of Interests

The authors declare that they have no known competing financial interests or personal relationships that could have appeared to influence the work reported in this paper.

## Notes

### Competing Interest Statement

The authors have declared no competing interest.

## References

Allred, B.W., Bestelmeyer, B.T., Boyd, C.S., Brown, C., Davies, K.W., Duniway, M.C., Ellsworth, L.M., Erickson, T.A., Fuhlendorf, S.D., Griffiths, T.V., Jansen, V., Jones, M.O., Karl, J., Knight, A., Maestas, J.D., Maynard, J.J., McCord, S.E., Naugle, D.E., Starns, H.D., Twidwell, D., and Uden, D.R. (2021). Improving Landsat predictions of rangeland fractional cover with multitask learning and uncertainty: Methods in Ecology and Evolution 12:5 841–849. 10.1111/2041-210X.13564.

Baruch-Mordo, S., Evans, J. S., Severson, J. P., Naugle, D. E., Maestas, J. D., Kiesecker, J. M., … & Reese, K. P. (2013). Saving sage-grouse from the trees: a proactive solution to reducing a key threat to a candidate species. Biological Conservation, 167, 233–241.

Bedrosian, G., Doherty, K. E., Martin, B. H., Theobald, D. M., Morford, S. L., Smith, J. T., Evans, J. S., Donnelly, J. P., Kumar, A. V., Guinotte, J., Heller, M. M. and D.E. Naugle. 2024. Cows, not plows: Using cropland conversion risk to scale-up averted loss of core sagebrush rangelands. Rangeland Ecology and Management: In Review.

Belton, V., Stewart, T.J. (2002). Implementation of MCDA: Practical Issues and Insights. In: Multiple Criteria Decision Analysis. Springer, Boston, MA. 10.1007/978-1-4615-1495-4_9

BLM: Bureau of Land Management. (2023). BLM National SMA Surface Management Agency Area Polygons. Bureau of Land Management, Denver, CO. https://gbp-blm-egis.hub.arcgis.com/datasets/blm-national-sma-surface-management-agency-area-polygons/about. Accessed January 3, 2024.

Boone, J. D., Ammon, E., & Johnson, K. (2018). Long-term declines in the Pinyon Jay and management implications for piñon–juniper woodlands. Trends and traditions: avifaunal change in western North America. Studies of Western Birds, 3, 190–197.

Boyd, C. S., Creutzburg, M. K., Kumar, A. V., Smith, J. T., Bradford, J. B., Cahill, M., Copeland, S., Doherty, K. E., Duquette, C., Garner, L., Holdrege, M. C., Mealor, B. A. and W. D. Sparklin. 2024. A strategic and science-based framework for management of invasive annual grasses in the sagebrush biome. Rangeland Ecology and Management: In Review.

Bybee, J., Roundy, B. A., Young, K. R., Hulet, A., Roundy, D. B., Crook, L., … & Cline, N. L. (2016). Vegetation response to piñon and juniper tree shredding. Rangeland Ecology & Management, 69(3), 224–234.

Caha J. and Drážná A. (2019). FuzzyAHP package for R (ver.0.9.5). R package version 0.9.5, http://github.com/JanCaha/FuzzyAHP/.

Chambers, J. C., Maestas, J. D., Pyke, D. A., Boyd, C. S., Pellant, M., & Wuenschel, A. (2017). Using resilience and resistance concepts to manage persistent threats to sagebrush ecosystems and greater sage-grouse. Rangeland Ecology & Management, 70(2), 149–164.

Coates, P. S., Prochazka, B., Webster, S. C., Weise, C. L., Aldridge, C. L., O’Donnell, M. S., Wiechman, L., Doherty, K. E. and J. C. Tull. Assessing performance of cooperative conservation actions on population growth of greater sage-grouse (Centrocercus urophasianus). Rangeland Ecology and Management: In Review.

Crist, M. R., Cross, T. B., Doherty, K. E., Olszewski, J. H. and K. C. Short. Will it burn? Characterizing wildfire risk for the Sagebrush Conservation Design. Rangeland Ecology and Management, In Review.

Daly, C., Halbleib, M., Smith, J. I., Gibson, W. P., Doggett, M. K., Taylor, G. H., … & Pasteris, P. P. (2008). Physiographically sensitive mapping of climatological temperature and precipitation across the conterminous United States. International Journal of Climatology: a Journal of the Royal Meteorological Society, 28(15), 2031–2064.

Doherty, K., Theobald, D. M., Bradford, J. B., Wiechman, L. A., Bedrosian, G., Boyd, C. S., … & Zeller, K. A. (2022). A sagebrush conservation design to proactively restore America’s sagebrush biome (No. 2022-1081). US Geological Survey.

Doherty, K.E., T. Remington, D.E. Naugle, J.D. Maestas, C.S. Boyd, L.A. Wiechman, J.B. Bradford, and G. Bedrosian. 2024. Sagebrush Conservation Design Phase 2: Implementing hope while managing change. Rangeland Ecology and Management: In Review.

Donnelly, J.P., Tack, J.D., Doherty, K.E., Naugle, D.E., Allred, B.W., Dreitz, V.J., 2017. Extending conifer removal and landscape protection strategies from sage-grouse to songbirds, a range-wide assessment. Rangel Ecol Manag 70, 95–105.

Fick, S. E., Nauman, T. W., Brungard, C. C., & Duniway, M. C. (2022). What determines the effectiveness of Pinyon-Juniper clearing treatments? Evidence from the remote sensing archive and counter-factual scenarios. Forest Ecology and Management, 505, 119879.

Filippelli, S. K., M. J. Falkowski, A. T. Hudak, P. A. Fekety, J. C. Vogeler, A. H. Khalyani, B. M. Rau, and E. K. Strand. (2020). “Monitoring Pinyon-Juniper Cover and Aboveground Biomass across the Great Basin.” Environmental Research Letters 15: 025004.

Flake, S. W., & Weisberg, P. J. (2019). Fine-scale stand structure mediates drought-induced tree mortality in pinyon–juniper woodlands. Ecological Applications, 29(2), e01831.

Holdrege, M. C., Palmquist, K. A., Schlaepfer, D. R., Lauenroth, W. K., Boyd, C. S., Creutzburg, M. K., Crist, M. R., Doherty, K. E., Remington, T. E., Tull, J. C., Wiechman, L. A. and J. B. Bradford. Climate change amplifies ongoing declines of sagebrush ecological integrity. Rangeland Ecology and Management: In Review.

Homer, C., Dewitz, J., Yang, L., Jin, S., Danielson, P., Xian, G., … & Megown, K. (2015). Completion of the 2011 National Land Cover Database for the conterminous United States–representing a decade of land cover change information. Photogrammetric Engineering & Remote Sensing, 81(5), 345–354.

Jarnevich, C., P. Engelstad, J. LaRoe, B. Hays, T. Hogan, J. Jirak, I. Pearse, J. Prevéy, J. Sieracki, A. Simpson, J. Wenick, N. Young, and H. R. Sofaer. 2023. Invaders at the doorstep: Using species distribution modeling to enhance invasive plant watch lists. Ecological Informatics, 75:101997.

Jarnevich, C.S., J. LaRoe, P. Engelstad, B. Hays, G. Henderson, D. Williams, K. Shadwell, I.S. Pearse, J.S. Prevey, and H.R. Sofaer. 2023. INHABIT species potential distribution across the contiguous United States (ver. 3.0, February 2023): U.S. Geological Survey data release.

Jeffries, M.I., and Finn, S.P. (2019). The sagebrush biome range extent, as derived from classified Landsat imagery: U.S. Geological Survey data release, accessed March 3, 2023, at 10.5066/P950H8HS

Kleinhesselink, A. R., Kachergis, E. J., McCord, S. E., Shirley, J., Hupp, N. R., Walker, J., … & Naugle, D. E. (2023). Long-Term Trends in Vegetation on Bureau of Land Management Rangelands in the Western United States. Rangeland Ecology & Management, 87, 1–12.

Kottek, M., Grieser, J., Beck, C., Rudolf, B., & Rubel, F. (2006). World map of the Köppen-Geiger climate classification updated. Meteorology Zeitschrift 15 (2006), 259–263.

Kumar, A.V.., Tack, J.D., Doherty, K. E., Smith, J.T.., Ross, B. E., and Bedrosian, G. 2024. Defend and grow the core for birds: How a biome-wide sagebrush conservation strategy benefits imperiled rangeland birds. Rangeland Ecology and Management: In Review.

Maestas, J., Jones, M., Pastick, N.J., Rigge, M.B., Wylie, B.K., Garner, L., Crist, M., Homer, C., Boyte, S., and Witacre, B. (2020). Annual herbaceous cover across rangelands of the sagebrush biome: U.S. Geological Survey data release, accessed March 1, 2023, at 10.5066/P9VL3LD5

Maestas, J.D., Naugle, D.E., Chambers, J.C., Tack, J.D., Boyd, C.S., and Tague, J.M., 2021, Conifer expansion, chap. M of Remington, T.E., Deibert, P.A., Hanser, S.E., Davis, D.M., Robb, L.A., and Welty, J.L., eds., Sagebrush conservation strategy—Challenges to sagebrush conservation: U.S. Geological Survey Open-File Report 2020–1125, p. 139–152, accessed September 29, 2023, at 10.3133/ofr20201125.

Marttunen, M., Lienert, J., & Belton, V. (2017). Structuring problems for Multi-Criteria Decision Analysis in practice: A literature review of method combinations. European journal of operational research, 263(1), 1–17.

Miller, R. F., J. C. Chambers, L. Evers, C. J. Williams, K. A. Snyder, B. A. Roundy, and F. B. Pierson. (2019). The Ecology, History, Ecohydrology, and Management of Pinyon and Juniper Woodlands in the Great Basin and Northern Colorado Plateau of the Western United States. General Technical Report. Fort Collins, CO: Rocky Mountain Research Station, USDA Forest Service.

Miller, R. F., Tausch, R. J., McArthur, E. D., Johnson, D. D., & Sanderson, S. C. (2008). Age Structure and Expansion of Piñon-juniper Woodlands: A Regional Perspective in the Intermountain West (pp. 1–15). Fort Collins, CO, USA: Rocky Mountain Research Station.

Miller, R.F.; Bates, J.D.; Svejcar, T.J.; [et al.] E. Eddleman. 2005. Biology, ecology, and management of western juniper (Juniperus occidentalis). Tech. Bul. 152. Corvallis, OR: Oregon State University Agricultural Experiment Station.

Morford, S.L., Allred, B.W., Jensen, E.R., Maestas, J.D., Mueller, K.R., Pacholski, C.L., Smith, J.T., Tack, J.D., Tackett, K.N. and Naugle, D.E. (2024), Mapping tree cover expansion in Montana, U.S.A. rangelands using high-resolution historical aerial imagery. Remote Sensing in Ecology and Conservation. 10.1002/rse2.357

Mozelewski, T., Freeman, P., Doherty, K. E., Kumar, A.V., Naugle, D. E., Morford, S. L., Kachergis, E. J., McCord, S. E., Jeffries, M., Pilliod, D. S. and L. A. Wiechman. 2024. State of the sagebrush: Conservation influences on the future of the biome. Rangeland Ecology and Management: In Review.

Naugle, D. E., Morford, S. L., Smith, J. T., Mueller, K. R., Griffiths, T. and T. Heater. Outcomes of spatial targeting in sagebrush country: A retrospective look at the NRCS-led Sage Grouse Initiative. Rangeland Ecology and Management: In Review.

Noel, A. R., Shriver, R. K., Crausbay, S. D., & Bradford, J. B. (2023). Where can managers effectively resist climate-driven ecological transformation in pinyon–juniper woodlands of the US Southwest?. Global Change Biology, 29((15), 4327–4341.

NRCS: Natural Resources Conservation Service. (2021). A framework for conservation action in the sagebrush biome—Working Lands for Wildlife: Washington, D.C., U.S. Department of Agriculture, Natural Resources Conservation Service, accessed December 12, 2023, at https://wlfw.rangelands.app.

Olsen, A. C., Severson, J. P., Maestas, J. D., Naugle, D. E., Smith, J. T., Tack, J. D., … & Hagen, C. A. (2021). Reversing tree expansion in sagebrush steppe yields population-level benefit for imperiled grouse. Ecosphere, 12(6), e03551.

R Core Team. (2023). R: A language and environment for statistical computing. R Foundation for Statistical Computing, Vienna, Austria. URL https://www.R-project.org/.

Reinhardt, J. R., Filippelli, S., Falkowski, M., Allred, B., Maestas, J. D., Carlson, J. C., & Naugle, D. E. (2020). Quantifying pinyon-juniper reduction within North America’s sagebrush ecosystem. Rangeland Ecology & Management, 73(3), 420–432.

Reinhardt, J. R., Naugle, D. E., Maestas, J. D., Allred, B., Evans, J., & Falkowski, M. (2017). Next-generation restoration for sage-grouse: a framework for visualizing local conifer cuts within a landscape context. Ecosphere, 8(7), e01888.

Reinhardt, J. R., Tack, J. D., Maestas, J. D., Naugle, D. E., Falkowski, M. J., & Doherty, K. E. (2023). Optimizing targeting of pinyon-juniper management for sagebrush birds of conservation concern while avoiding imperiled pinyon jay. Rangeland Ecology & Management, 88, 62–69.

Romme, W. H., C. D. Allen, J. D. Bailey, W. L. Baker, B. T. Bestelmeyer, P. M. Brown, K. S. Eisenhart, et al. (2009). “Historical and Modern Disturbance Regimes, Stand Structures, and Landscape Dynamics in Piñon–Juniper Vegetation of the Western United States.” Rangeland Ecology & Management 62: 203–22.

Roundy, B. A., Miller, R. F., Tausch, R. J., Young, K., Hulet, A., Rau, B., … & Eggett, D. (2014). Understory cover responses to piñon–juniper treatments across tree dominance gradients in the Great Basin. Rangeland Ecology & Management, 67(5), 482–494.

Saaty, R. W. (1987). The analytic hierarchy process—what it is and how it is used. Mathematical modelling, 9(3-5), 161–176.

Schultz, E. L., Filippelli, S. K., Vogeler, J. C., & Shriver, R. K. (2023). Density-dependent dynamics help explain the simultaneous expansion and decline of woodlands in the western US. Forest Ecology and Management, 546, 121359.

Schultz, E. L., Hülsmann, L., Pillet, M. D., Hartig, F., Breshears, D. D., Record, S., … & Evans, M. E. (2022). Climate-driven, but dynamic and complex? A reconciliation of competing hypotheses for species’ distributions. Ecology letters, 25(1), 38–51.

Severson, J. P., Hagen, C. A., Maestas, J. D., Naugle, D. E., Forbes, J. T., & Reese, K. P. (2017). Short-term response of sage-grouse nesting to conifer removal in the northern Great Basin. Rangeland Ecology & Management, 70(1), 50–58.

Shriver, R. K., Yackulic, C. B., Bell, D. M., & Bradford, J. B. (2022). Dry forest decline is driven by both declining recruitment and increasing mortality in response to warm, dry conditions. Global Ecology and Biogeography, 31(11), 2259–2269.

Smith, J.T., White, C., Morford, S. L., Maestas, J. D., Kleinhesselink, A. R. and D.E. Naugle. Monitoring Rangeland vegetation response to conifer removal in southeastern Idaho via Satellite Imagery. Rangeland Ecology and Management: In Review.

Snyder, K. A., Evers, L., Chambers, J. C., Dunham, J., Bradford, J. B., & Loik, M. E. (2019). Effects of changing climate on the hydrological cycle in cold desert ecosystems of the Great Basin and Columbia Plateau. Rangeland Ecology & Management, 72(1), 1–12.

Tack, J. D., Smith, J. T., Doherty, K. E., Donnelly, P. J., Maestas, J. D., Allred, B. W., … & Naugle, D. E. (2023). Regional Context for Balancing Sagebrush-and Woodland-Dependent Songbird Needs with Targeted Pinyon-Juniper Management. Rangeland Ecology & Management, 88, 182–191.

Theobald, D. M., Kumar, A. V., Doherty, K. E., Zeller, K. A., Cross, T. and S. Finn. Anchoring sagebrush conservation to core landscapes by understanding the decline of sagebrush ecosystem connectivity from 2001-2021. Rangeland Ecology and Management: In Review.

USGS GAP: U.S. Geological Survey Gap Analysis Project. (2016). GAP/LANDFIRE National Terrestrial Ecosystems 2011: U.S. Geological Survey data release, accessed April 17, 2023, at 10.5066/F7ZS2TM0.

Vander Wall, S. B. (2023). Seed Dispersal in Pines (Pinus). The Botanical Review, 1–33.

Williams, D.A., C.S. Jarnevich, P. Engelstad, G. Henderson, K. Shadwell, I.S. Pearse, and J.S. Prevey. 2023. Potential distribution of Japanese brome (Bromus japonicus) across the contiguous United States (October 2023): U.S. Geological Survey data release

Wollstein, K., Johnson, D. and C. Boyd. Operationalizing strategic conservation: A multi-level framework to identify opportunities and actions. Rangeland Ecology and Management: In Review.

Zeller, K. A., Cushman, S. A., Van Lanen, N. J., Boone, J. D., & Ammon, E. (2021). Targeting conifer removal to create an even playing field for birds in the Great Basin. Biological Conservation, 257, 109130.

